# Transcriptomic Profile Analysis of Brain Inferior Colliculus Following Acute Hydrogen Sulfide Exposure

**DOI:** 10.1101/816710

**Authors:** Dong-Suk Kim, Poojya Anantharam, Piyush Padhi, Daniel R Thedens, Ganwu Li, Ebony Gilbreath, Wilson K. Rumbeiha

## Abstract

Hydrogen sulfide (H_2_S) is a gaseous molecule found naturally in the environment, and as an industrial byproduct, and is known to cause acute death and induces long-term neurological disorders following acute high dose exposures. Currently, there is no drug approved for treatment of acute H_2_S-induced neurotoxicity and/or neurological sequelae. Lack of a deep understanding of pathogenesis of H_2_S-induced neurotoxicity has delayed the development of appropriate therapeutic drugs that target H_2_S-induced neuropathology. RNA sequencing analysis was performed to elucidate the cellular and molecular mechanisms of H_2_S-induced neurodegeneration, and to identify key molecular elements and pathways that contribute to H_2_S-induced neurotoxicity. C57BL/6J mice were exposed by whole body inhalation to 700 ppm of H_2_S for either one day, two consecutive days or 4 consecutive days. Magnetic resonance imaging (MRI) scan analyses showed H_2_S exposure induced lesions in the inferior colliculus (IC) and thalamus (TH). This mechanistic study focused on the IC. RNA Sequencing analysis revealed that mice exposed once, twice, or 4 times had 283, 193 and 296 differentially expressed genes (DEG), respectively (q-value < 0.05, fold-change > 1.5). Hydrogen sulfide exposure modulated multiple biological pathways including unfolded protein response, neurotransmitters, oxidative stress, hypoxia, calcium signaling, and inflammatory response in the IC. Hydrogen sulfide exposure activated PI3K/Akt and MAPK signaling pathways. Pro-inflammatory cytokines were shown to be potential initiators of the modulated signaling pathways following H_2_S exposure. Furthermore, microglia were shown to release IL-18 and astrocytes released both IL-1β and IL-18 in response to H_2_S. This transcriptomic analysis data revealed complex signaling pathways involved in H_2_S-induced neurotoxicity and may provide important associated mechanistic insights.

**Highlights:**

- Transcriptomic profiling analyses following acute exposure to H_2_S were performed
- Multiple signaling pathways were dysregulated following H_2_S exposure
- PI3K/Akt and MAPK signaling pathways were activated after H_2_S exposure
- MRI scan analysis revealed lesions in the IC and TH following H_2_S exposure
- Acute H_2_S exposure induced a neuroinflammatory response

## 1. Introduction

Hydrogen sulfide (H_2_S) is a flammable and colorless potent toxic gas with a “rotten egg” odor that has generally been considered as an environmental hazard (Chou et al. 2016). However, recently it has been shown that H_2_S is endogenously produced in low concentrations in multiple organs including heart, brain, kidney, and aorta (Olas 2014). In low physiological concentrations H_2_S plays important roles including acting as a gasotransmitter, regulating vascular tone (Lavu et al. 2011), serving as an anti-inflammatory compound (Lavu et al. 2011), and as a scavenger of reactive oxygen species (ROS) (Lavu et al. 2011). It also plays a beneficial role in several diseases including cancer (Lee et al. 2011) and Parkinson’s disease (Hu et al. 2010). Hydrogen sulfide has also been shown to be involved in multiple signaling pathways including protein kinase A, mitogen and receptor tyrosine kinases, oxidative stress, calcium channels and neurotransmission, among others (Tan et al. 2010).

Acute exposure to high concentrations of H_2_S induces concentration-, time-and dose-dependent toxicity in humans and animals (Anantharam et al. 2017; Board 2004; Rumbeiha et al. 2016). It is the second most common cause of acute toxic gas induced mortality, after carbon monoxide (Greenberg and Hamilton 1998). Acute exposure to 30 ppm H_2_S induces olfactory fatigue and malfunction of the nose. At concentrations higher than 500 ppm H_2_S induces headache, dizziness and respiratory failure. Additional effects of H_2_S concentrations > 500 ppm include respiratory paralysis, seizures, and loss of consciousness (knock-down). A brief exposure to H_2_S > 1000 ppm can be instantly fatal. Environmental H_2_S is released naturally from volcanoes, sulfur springs, stagnant water bodies and geothermally active areas. Hydrogen sulfide can also be produced by human activities such as desulfurization processes in oil and gas industries, paper mills, manure treatment facilities, and in water purification facilities. It is a toxic industrial raw chemical material. Hydrogen sulfide can easily be generated from mixing simple raw materials and has been used for suicide purposes (Sams et al. 2013). In addition, H_2_S can potentially be used for nefarious activities by terrorists (National Institute of Allergy and Infectious Diseases 2007; Ng et al. 2019); therefore, potential misuse of this potent toxicant by terrorists is concerning and a major risk to victims and first responders (Ng et al. 2019; Sams et al. 2013). Survivors of acute exposure to high concentrations of H_2_S may manifest one or more neurological deficits including slurred speech, memory loss, dizziness, sleep disturbances, depression, delirium, altered psychological states, spasms, and motor dysfunction (Board 2004; Chou et al. 2016; Guidotti 2010; Guidotti 2015; Kilburn 2003; Matsuo et al. 1979; Parra et al. 1991; Rumbeiha et al. 2016; Tvedt et al. 1991a; Tvedt et al. 1991b; Wasch et al. 1989). There are more than 1,000 annual reports related to severe H_2_S exposure in the Unites States (U.S. Department of Health and Human Services 2014). Despite the high mortality and morbidity of acute H_2_S poisoning, there is no suitable antidote available for field use to treat civilian casualties and to prevent development of incapacitating neurological sequelae.

The exact molecular and cellular mechanisms behind H_2_S-induced neurological disorders are poorly understood and this is the main reason responsible for delayed development of drugs for treatment of H_2_S-induced acute toxicity, though a few toxic mechanisms have been suggested. For example, cytochrome c dependent apoptosis with DNA damage was previously reported to play a role in H_2_S-induced neurotoxicity (Kurokawa et al. 2011). Glutamate-induced cytotoxicity was also reported to be involved in H_2_S exposure-induced neurotoxicity (Cheung et al. 2007; Garcia-Bereguiain et al. 2008). Reduction of glutathione (GSH), an antioxidant, and generation of ROS have also been shown to play a role *in vitro* (Truong et al. 2006). Most of these studies were done *in vitro* and have not been confirmed in *in vivo* animal models. There is still a wide knowledge gap in understanding the mechanisms of H_2_S-induced neurotoxicity. Closing this gap is critical for the development of novel therapeutic antidotes.

In this study, we used a relevant inhalation mouse model of H_2_S-induced neurotoxicity developed in-house to study molecular mechanisms of acute H_2_S-induced neurotoxicity using transcriptomic analysis. This mouse model manifests motor and behavioral deficits, respiratory depression, seizures, and knock-down, recapitulating the human condition (Anantharam et al. 2017). In this study, we have shown that magnetic resonance imaging (MRI) analysis from live mice exposed to H_2_S revealed neurodegeneration in the inferior colliculus (IC) and in the thalamus (TH). We focused on the neurodegeneration in the IC, the most severely affected region, and showed that transcriptomic analyses of the IC of mice acutely exposed to H_2_S had multiple modulated biological pathways and potential upstream regulators. Among these, the PI3K/Akt and MAPK signaling pathways were activated following acute H_2_S exposure. We showed that microglia and astrocytes released the pro-inflammatory cytokines, IL-1β and IL-18. This data provides valuable information on molecular targets which may play an important role in the development of novel antidotes or therapies for acute H_2_S-induced neurotoxicity.

## 2. Materials and Methods

### 2.1 Chemicals

Hydrogen sulfide gas [CAS 7783-06-4] was purchased from Airgas (Radnor, PA). RNeasy mini kit was purchased from Qiagen (Germantown, MD). High Capacity cDNA RT kit were purchased from ThermoFisher Scientific (Waltham, MA). RT² SYBR Green ROX qPCR Mastermix and primers for Gapdh were purchased from Qiagen (Valencia, CA).

### 2.2 Animals

All animal studies were approved by the Iowa State University Institutional Animal Care and Use Committee (IACUC). Animals were cared for in accordance with IACUC guidelines. Mice were purchased from Jackson Laboratories (Bar Harbor, ME). Seven-to eight-week-old male C57 BL/6J mice were housed at room temperature (RT) of 20 −22 °C under a 12-h light cycle, and a relative humidity of 35 – 50 %. Protein Rodent maintenance diet (Teklad HSD Inc, WI, US) and water were provided *ad libitum*. All mice were acclimated to breathing air for 1 week before H_2_S exposure. We chose to use the C57BL/6J mouse model because it is widely used for neurodegenerative research. This model has effectively recapitulated H_2_S-induced neuropathology in our hands (Anantharam et al. 2018a; Anantharam et al. 2017). In this study, we used male mice only because, as previously reported, male mice were more sensitive than female mice (Anantharam et al. 2018a; Anantharam et al. 2017).

### 2.3 Transcriptomic analysis

#### 2.3.1 Exposure paradigm

For transcriptomic analysis, mice were exposed to H_2_S either once, twice or 4 times. Mice exposed once reflect the typical human accidental exposure scenario. Mice exposed 2 or 4 times were used for comparison to the typical one-time exposure. Mice were exposed by whole body inhalation either to normal breathing air from a tank (Control) or to 700ppm H_2_S for 40 min on day 1 for mice which received one exposure. For mice which received 2 exposures, they were exposed to 700ppm H_2_S for 40 min on day 1 and for 15 min on day 2. For mice which received 4 exposures they received 700ppm H_2_S for 40 min on day 1 and for 15 mins per day on subsequent days up to day 4 (Fig. 1). Control group mice were exposed to breathing air daily for 4 days. We chose this H_2_S exposure level of 700 ppm because we are mimicking high dose exposures as would be encountered during terrorism incidents or accidents in which concentrations of H_2_S exceed 500 ppm. The duration of exposure in field accidents can vary depending on how quickly first responders get to the site, terrain, etc. For our studies, exposure duration was based on preliminary dose-response and time course studies (Anantharam et al. 2017). Following rapid decapitation, brains were immediately removed from the skull. The IC of the four mice in each group were microdissected on ice and immediately flash frozen using liquid nitrogen and stored at −80°C until further use.

**Figure 1.**
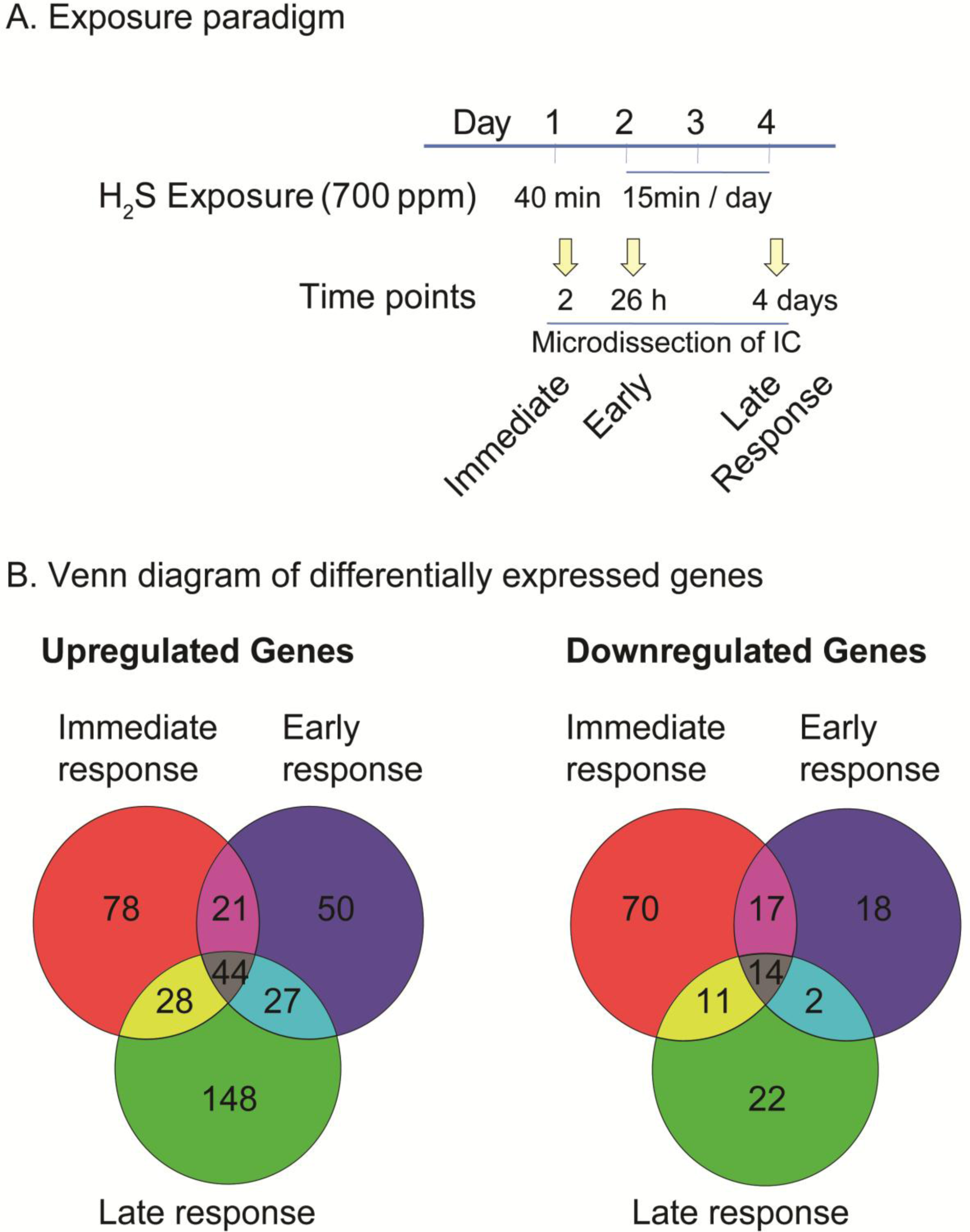
Hydrogen sulfide exposure paradigm and Venn diagram of differentially expressed genes (DEGs) after acute exposure to H_2_S in the IC. C57 black mice were exposed via whole body exposure to 700 ppm H_2_S. A scheme of exposure paradigm was shown (A). The microdissected IC was used for transcriptomic analysis. The numbers of DEGs at immediate, early, and late response to H_2_S exposure in the IC are shown in Venn diagrams (B).

#### 2.3.2 RNA isolation

Total RNA was extracted from frozen tissues using the RNeasy® Plus Mini kit with treatment of DNase I according to the manufacturer’s protocol (Qiagen, Germantown, MD). The integrity of RNA samples were examined by running on agarose gel and using bioanalyzer.

#### 2.3.3 Transcriptome profiling with RNA-Seq

Construction of library of samples was followed by TruSeq® RNA Sample Preparation v2 Guide (Illumina). Briefly, poly (A)-containing mRNA of samples were used for first-strand cDNA synthesis using Super Script II (Invitrogen). RNA Index adapter were ligated to the cDNA products, followed by several rounds of PCR amplification. The resulting libraries of samples were sequenced using HiSeq 2000 System (Illumina). The library construction and RNA-Seq were performed at BGI America (Cambridge, MA).

#### 2.3.4 Transcriptome analysis

Transcriptomic profiling was processed following Tuxedo protocol (nature paper) with slight modification. For the modifications, we used STAR software for mapping raw data to the mouse reference genome (GRCm38.p4 downloaded from https://www.gencodegenes.org/mouse/release_M11.html). STAR (version 2.5.1), samtools (version 0.1.19), Bowtie 2 (version 2.2.7), cufflinks (version 2.2.1), tophat 2 (version 2.1.0) were used for the analyses. Differentially expressed genes (DEG) were obtained by using cuffdiff, part of Tuxedo suite software. Ingenuity Pathway Analysis (Qiagen) was used to analyze gene ontology analyses including altered biological pathways and potential initiator. DEGs common in immediate, early, and late response were used to generate heat map by using R software (version 3. 3. 2, https://cran.r-project.org). Ingenuity Pathway Analysis software (Qiagen, Germantown, MD) was used for dysregulated biological pathway and potential upstream regulator analysis.

#### 2.3.5 Validation of gene expression via quantitative real-time RT-PCR

Total RNAs were extracted from frozen tissues using the RNeasy® Plus Mini kit and treated with DNase I (Invitrogen) according to the manufacturer’s protocol. cDNA was prepared using High-Capacity cDNA Reverse Transcription Kits from ThermoFisher Scientific (Waltham, MA). RT^2^ SYBR Green ROX qPCR Mastermix and primers for Gapdh were purchased from Qiagen (Valencia, CA). The threshold cycle (C_t_) was calculated from the instrument software, and fold change in gene expression was calculated using the ΔΔC_t_ method as described earlier (Kim et al. 2016). Primer sequences will be provided upon request.

### 2.4 Western blot analysis

For Western blot analysis mice were exposed to H_2_S once a day for up to 7 days. Mice were exposed to 700ppm H_2_S for 40 min on day 1 (single exposure) or to 700ppm H_2_S for 40 min on day 1 and for 15 min per day on subsequent days up to day 7 (2 to 7 exposures). Control group mice were exposed to breathing air from a tank daily for 7 days (Fig. 4). Following rapid decapitation, brains were immediately removed from the skull. The IC of three mice in each group were microdissected on ice and immediately flash frozen using liquid nitrogen and stored at −80°C until further use. Samples were lysed with modified RIPA lysis buffer 1% Triton X-100, 1 mM EDTA, 100 mM NaCl, 1 mM EGTA, 1 mM NaF, 20 mM Na_4_P_2_O_7_, 2 mM Na_3_VO_4_, 10% glycerol, 0.1% SDS, 0.5% deoxycholate, 50 mM Tris-HCl, and pH 7.4) via sonication, followed by centrifugation as described previously (Kim et al. 2016). Briefly, the samples containing equal amounts of proteins were loaded and fractionated in SDS-PAGE gel and transferred onto a nitrocellulose membrane overnight at 4°C. Membranes were blocked with 5% bovine serum albumin in TBS supplemented with 0.1 % Tween-20. Primary antibodies against specific proteins were incubated with the membrane overnight at 4°C.

The antibodies detecting phosphorylated proteins were purchased from Cell Signaling Technology (Cell Signaling Technology, Danvers, MA). Primary antibodies against PKA Cα, PI3K and pan-Akt were purchased from Cell signaling (Cell Signaling Technology, Danvers, MA). Primary antibodies against ATF2, and CREB-1 were purchased from Santa Cruz Biotechnology (Santa Cruz Biochnoloby, Dallas, Tx). After rinsing thoroughly in PBS supplemented with 0.1% Tween-20, the membrane was incubated with secondary antibodies. For the loading control, β-actin antibody was used. Immunoblot imaging was performed with an Odyssey Infrared Imaging system (LI-COR, Lincoln, NE). ImageJ software (National Institutes of Health, Bethesda, MD) was used to quantify Western blot bands of targeted proteins.

### 2.5 Magnetic resonance imaging analysis

For proof-of-principle, mice for MRI analysis were exposed by whole body inhalation either to normal breathing air from a tank (Control) or to 700ppm H_2_S for 40 min on day 1 and for 15 min per day on the subsequent 6 days (Fig. 6). Five mice in each group were transported to University of Iowa where MRI scanning was performed. MRI was performed using a Discovery MR901 7T small animal imager (GE Healthcare) at the University of Iowa, Iowa City, IA. Mice were anesthetized with isoflurane (3% in an induction followed by 1.5% via a nose cone for maintenance at 0.8 l/min flow rate in oxygen). Mice were kept warm during imaging. For T2 weighted imaging of the whole brain, a fast spin echo sequence was used (TR = 9100 ms, TE = 80 ms, matrix 256 × 192, number of slices = 40, field of view = 2.5 × 1.9 cm, slice thickness = 0.4 mm, 12 echoes, echo spacing = 8.9 ms, number of averages = 10, total acquisition time approximately 18 min). Brain lesion sizes were quantified using ImageJ software (NIH) by measuring the area of lesions.

### 2.6 Cell culture and quantitative RT-PCR for inflammatory response analysis

Mouse microglial cells, MMC, were kindly gifted from Dr. Anumantha Kanthasamy (Iowa State University, Ames, IA). MMC cells have been shown to mimic neonatal primary mouse microglia and to be a good mouse microglia model for neurotoxicity study (Sarkar et al. 2018b). U373 are established astrocytoid cells from human astrocytoma cell line and are suitable for neurotoxicity studies (Sarkar et al. 2018a). U373 and MMC were maintained in DMEM/F-12 media supplemented with 10% fetal bovine serum, 2 mM L-glutamine, 50 units of penicillin, and 50 μg/ml of streptomycin in a humidified incubator with 5% CO_2_ at 37°C as described previously (Kim et al. 2017). Cells were seeded to confluent at 50% next day. Cells were exposed to 1 mM of Sodium hydrosulfide (NaHS) [CAS 16721-80-5] for 6 h before harvesting total RNA. Total RNAs were used to produce cDNA for quantitative RT-PCR.

### 2.7 Statistical analysis

Data were analyzed using Prism 4.0 (GraphPad Software, San Diego, CA). Non-paired Student’s *t*-test was used when two groups were being compared. Differences were considered statistically significant at p-values <0.05. Data are represented as the mean ± S.E.M. of at least two separate experiments performed at least in triplicate.

## 3. Results

### 3.1 Acute exposure to H_2_S induced brain lesions in thalamus and inferior colliculus in C57 black mice

A previous study of ours revealed that H_2_S exposure induced overt necrosis most consistently in the IC (Anantharam et al. 2017). It is for this reason that we chose to study the IC in this experiment. In this study, we sought transcriptomic profiling analyses in the IC of H_2_S-exposed mice to search for key molecules involved in H_2_S-induced neurotoxicity at various time points as previously described (Fig. 1). Immediate response was measured at 2 h post a single H_2_S exposure. Early response was measured at 26 h post H_2_S exposure (mice were subjected to 2 acute H_2_S exposures 24 hrs apart and euthanized 2 h after the second exposure). The late response was measured following 4 days of acute H_2_S exposure. Mice exposed to H_2_S showed motor-behavioral deficits, respiratory depression, seizures and knock-down phenotype as previously reported ((Anantharam et al. 2017; Kim et al. 2018b)).

### 3.2 H_2_S exposure induced altered transcriptomic profiling in the IC

The IC was used for transcriptomic profiling analysis because it is one of the two most sensitive brain region following acute H_2_S exposure following the exposure paradigm used in this study. Transcriptomic profiling analyses of the IC were performed at 2 h, day 2, and day 4 after H_2_S exposure (see Fig. 1A) following Tuxedo protocol (Trapnell et al. 2012). The numbers of significantly up-regulated and down-regulated genes are shown in the Venn diagram (Fig 1B). 171, 142, and 247 genes were upregulated after H_2_S exposure, respectively at 2h, 26h, and 4 days, respectively. 112, 51, and 49 genes were downregulated following H_2_S exposure, at 2 h, 26 h, and 4 days, respectively. The genes that are up-regulated and down-regulated in common at all three time points are shown in the heatmap in Fig. 2.

**Figure 2.**
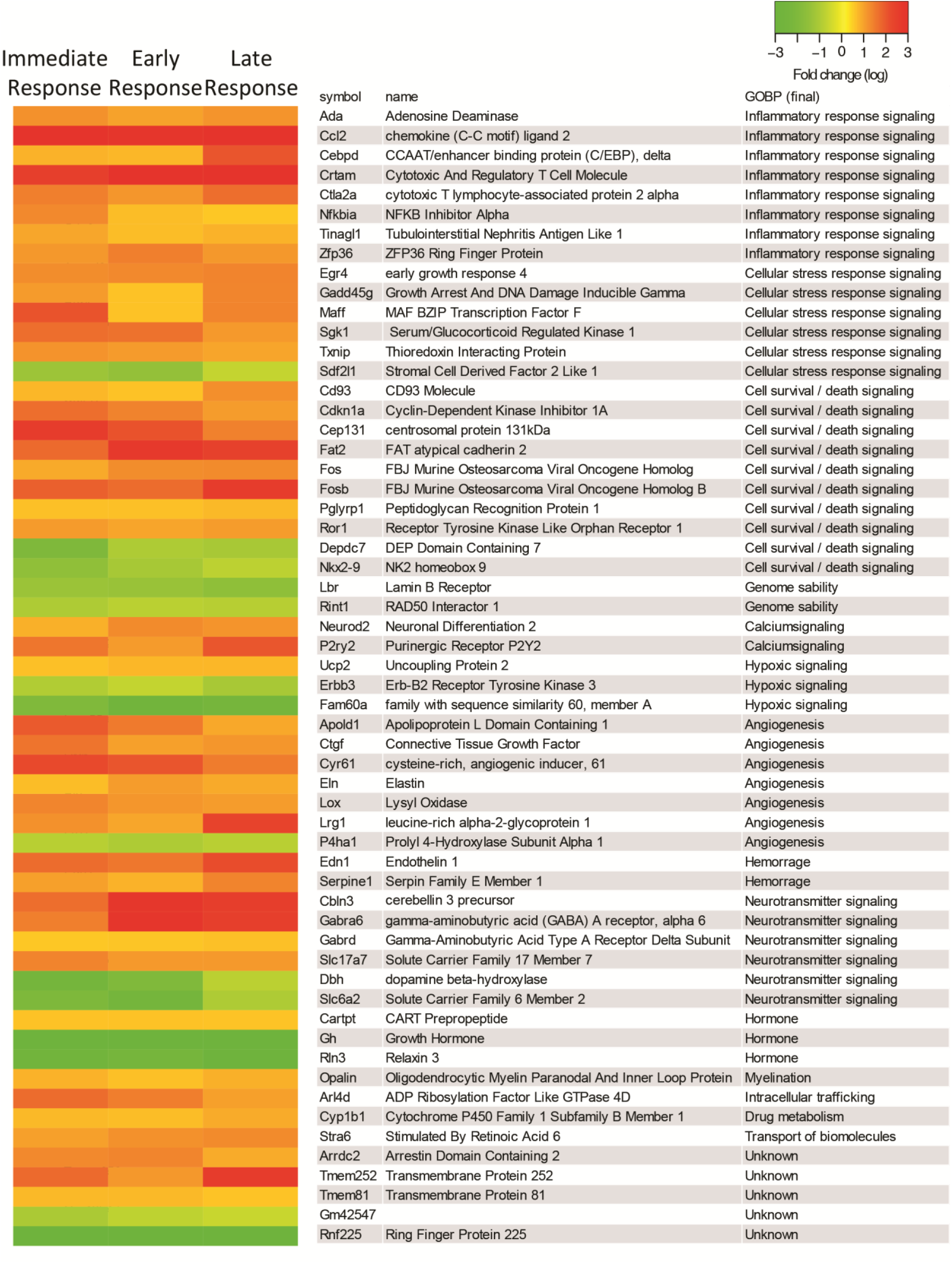
Heatmaps of DEGs in the IC after acute exposure to H_2_S. RNA-seq analysis of gene expression changes in the IC after H_2_S exposure. Expression of DEGs at immediate, early, and late response to H_2_S exposure are shown in heat map view. Genes with higher expression levels are shown in red, whereas genes with lower expression levels are shown in green. Gene names and gene ontology of biological pathways are shown in the right panel.

### 3.3 H_2_S exposure caused dysregulation of molecular biological pathways in the IC

Differentially expressed genes were used to analyze the dysregulated molecular biological pathways in the IC following H_2_S exposure. The top 10 dysregulated pathways are listed in table 1. Results show that a single H_2_S exposure and two acute H_2_S exposures shared many common altered pathways including unfolded protein response, coagulation system, and NRF2-mediated oxidative stress responses. A single H_2_S exposure by itself induced multiple dysregulated biological pathways (Table 1). In contrast, 4 days of repeated daily H_2_S exposures induced inflammation-related pathways (Table 1). The top 10 potential upstream regulators were also analyzed (table 2) for each time point. Note that factors related to initiation of the inflammatory response were found to be common throughout the study period, including 2 hrs following a single exposure (immediate response) (Table 1).

**Table 1.** The top 10 dysregulated biological pathways at immediate (2 h), early (26 h) and late (day 4) response in IC following acute H2S exposure. Dysregulated biological pathways were analyzed for immediate, early, and late response to H2S exposure. The top 10 dysregulated biological pathways were measured by statistical analyses and are listed. Degree of activating biological pathways is shown by the z-Score. Higher value of z-Score indicates activation of biological pathways, whereas lower value of z-Score for inactivation of biological pathways.

| Exposure | Pathway | pValue | z-Score |
| --- | --- | --- | --- |
| Immediate response | Unfolded protein response | 1.05E-06 |  |
| Immediate response | Catecholamine Biosynthesis | 6.72E-05 |  |
| Immediate response | Coagulation System | 2.48E-04 |  |
| Immediate response | GADD45 Signaling | 1.25E-03 |  |
| Immediate response | Dopamine Receptor Signaling | 3.73E-03 |  |
| Immediate response | NRF2-mediated Oxidative Stress Response | 4.51E-03 | 1.342 |
| Immediate response | GABA Receptor Signaling | 7.61E-03 | 1 |
| Immediate response | Death Receptor Signaling | 9.78E-03 | 0.378 |
| Immediate response | eNOS Signaling | 1.95E-02 | 0.707 |
| Immediate response | p53 Signaling | 2.51E-02 | 2 |
| Early response | Unfolded protein response | 2.05E-07 |  |
| Early response | NRF2-mediated Oxidative Stress Response | 8.26E-05 | 0.447 |
| Early response | Hypoxia Signaling in the Cardiovascular System | 8.94E-05 | 1.633 |
| Early response | Coagulation System | 2.44E-04 | -0.447 |
| Early response | Endoplasmic Reticulum Stress Pathway | 3.34E-04 |  |
| Early response | Acute Phase Response Signaling | 4.74E-04 | 2.646 |
| Early response | PI3K Signaling in B Lymphocytes | 5.08E-03 | 0.816 |
| Early response | Calcium Signaling | 9.02E-03 | -0.447 |
| Early response | IL-6 Signaling | 1.80E-02 | 0.816 |
| Early response | HMGB1 Signaling | 2.20E-02 | 1.342 |
| Late response | Agranulocyte Adhesion and Diapedesis | 1.55E-10 |  |
| Late response | Granulocyte Adhesion and Diapedesis | 1.08E-08 |  |
| Late response | Acute Phase Response Signaling | 1.67E-07 | 2.887 |
| Late response | HMGB1 Signaling | 2.00E-07 | 2.673 |
| Late response | IL-10 Signaling | 2.18E-07 |  |
| Late response | Coagulation System | 8.45E-06 | -1.134 |
| Late response | phagosome formation | 1.38E-05 |  |
| Late response | IL-6 Signaling | 1.01E-04 | 1.897 |
| Late response | JAK/Stat Signaling | 4.32E-04 | 0.707 |
| Late response | PPAR Signaling | 9.24E-04 | -2.828 |

**Table 2.**
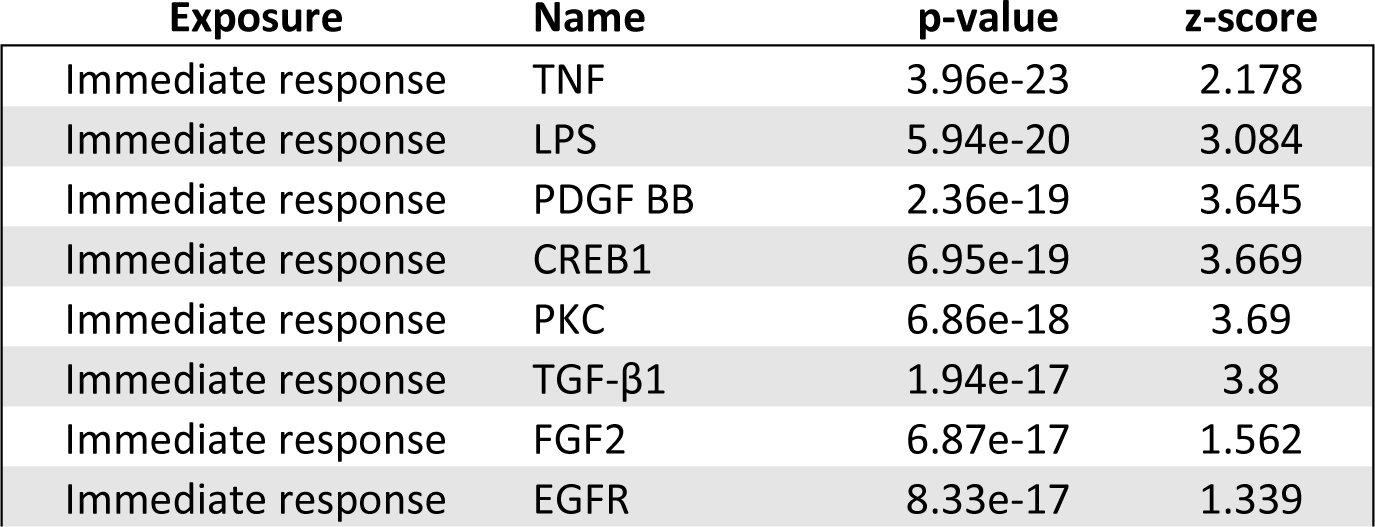
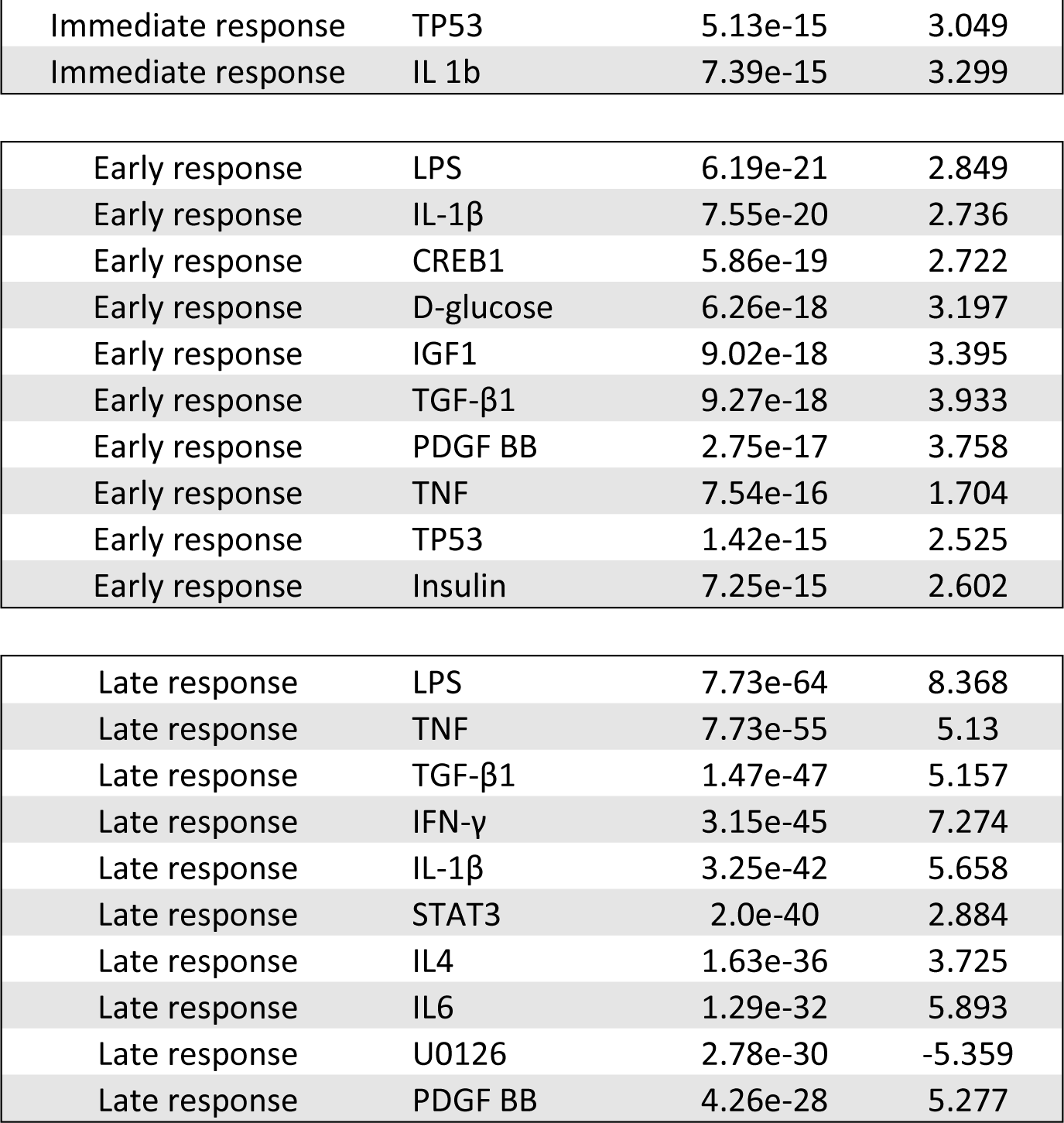
Potential upstream regulators in the IC responding to acute H2S exposure. Potential upstream regulators were analyzed at immediate (2 h), early (26 h), and late response (day 4) to acute H2S exposure. The top 10 potential upstream regulators were statistically analyzed and are listed. Degree of activating biological pathways is shown by z-Score. Higher value of z-Score indicates activation of potential upstream regulator, whereas lower value of z-Score for inactivation of potential upstream regulator.

### 3.4 Validation of gene expression profiles using quantitative real-time PCR

Quantitative real-time PCR of selected genes was performed to validate the gene expression profiles in the IC. Results show that genes that were significantly differentially expressed were shown to have similar trends with results of quantitative real-time PCR (Fig. 3). Selected genes that were differentially expressed in common throughout H_2_S exposure are shown in Fig. 3.

**Figure 3.**
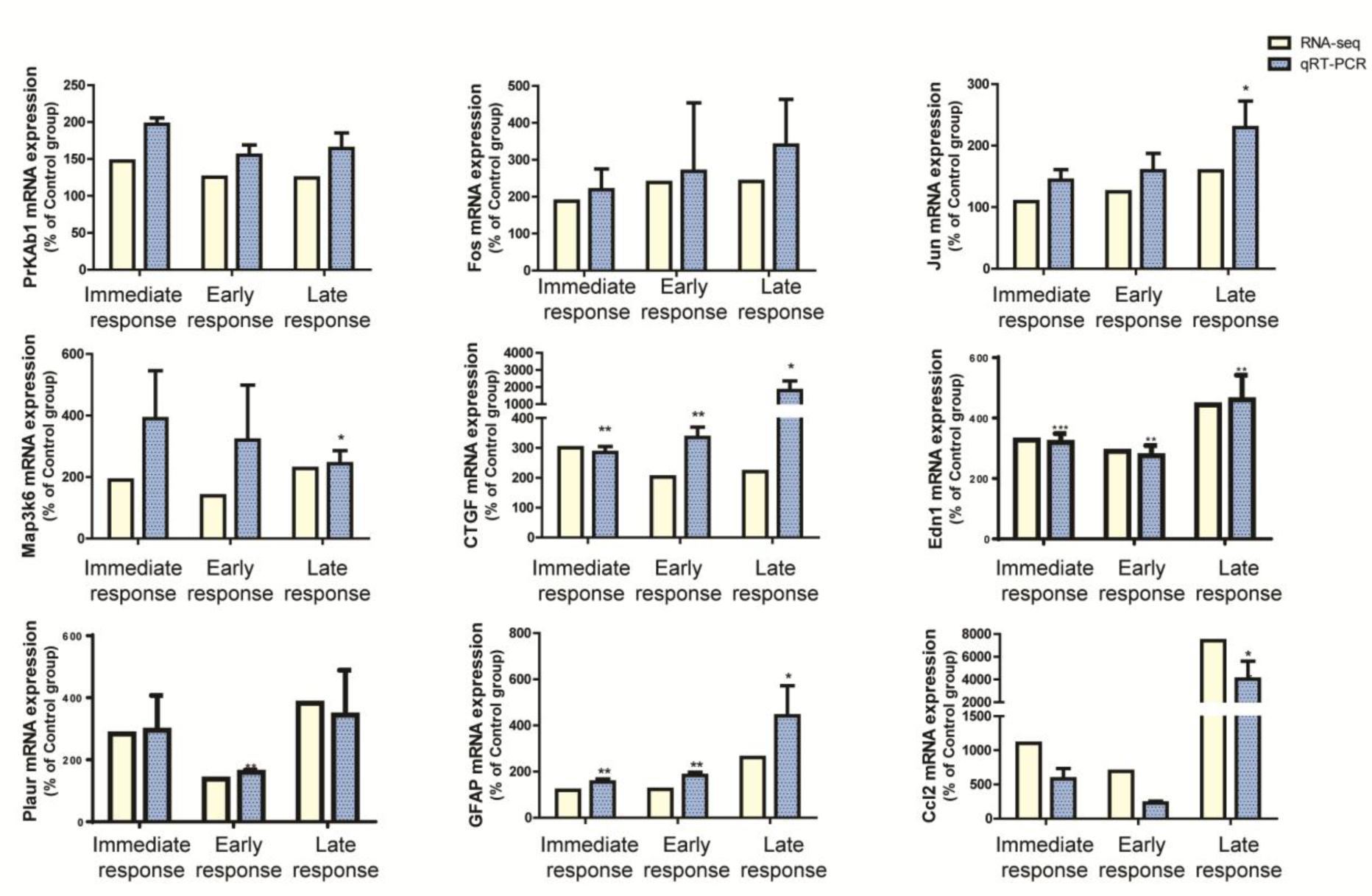
RNAseq analysis of the IC of H_2_S exposed mice were validated with quantitative real-time PCR. Expression profiles of RNAseq analyses in the IC were validated with the analyses of quantitative real-time PCR. Genes that were differentially expressed in common throughout H_2_S exposure were selected for this analysis. Expressions of genes were measured by quantitative real-time PCR (n>3) and compared to gene expression profiles of RNAseq analyses in the IC. Values of quantitative real-time PCR were normalized to Gapdh expression. Significant difference were determined by t-test by comparison to the breathing air control group and were indicated with *(p<0.05) or **(p<0.01).

### 3.5 H_2_S exposure induces activation of PI3K/Akt signaling pathway

Akt plays an important role in cell proliferation, cell death, and many other functions (Los et al. 2009) and mediates multiple signaling pathways including calcium signaling, JNK pathway, GABA signaling, mTOR signaling and cell death signaling which were all shown to be dysregulated by acute H_2_S exposure (Table 1). Phosphorylation of Akt at threonine 308 and at Serine 473 was increased by more than two fold at 2h following a single acute H_2_S exposure and 26 h post H_2_S exposure [2 h following a second acute H_2_S exposure], indicating activation of Akt (Fig. 4). Expression of phosphoinositide 3-kinases (PI3K) also followed a similar trend following H_2_S exposure (Fig. 4). These results show that exposure to H_2_S may activate the PI3K/Akt signaling pathway. Phosphorylation of cAMP-dependent protein kinase (PKA) shares substrate specificity with Akt. C subunit of PKA at Thr 197 was increased in the early phase of H_2_S exposure (Fig 4). Akt activates many substrates downstream in the signaling cascade. P70 S6 kinase, one of downstream substrates of Akt signaling, was phosphorylated at Thr 389 at 2h following a single acute H_2_S exposure.

**Figure 4.**
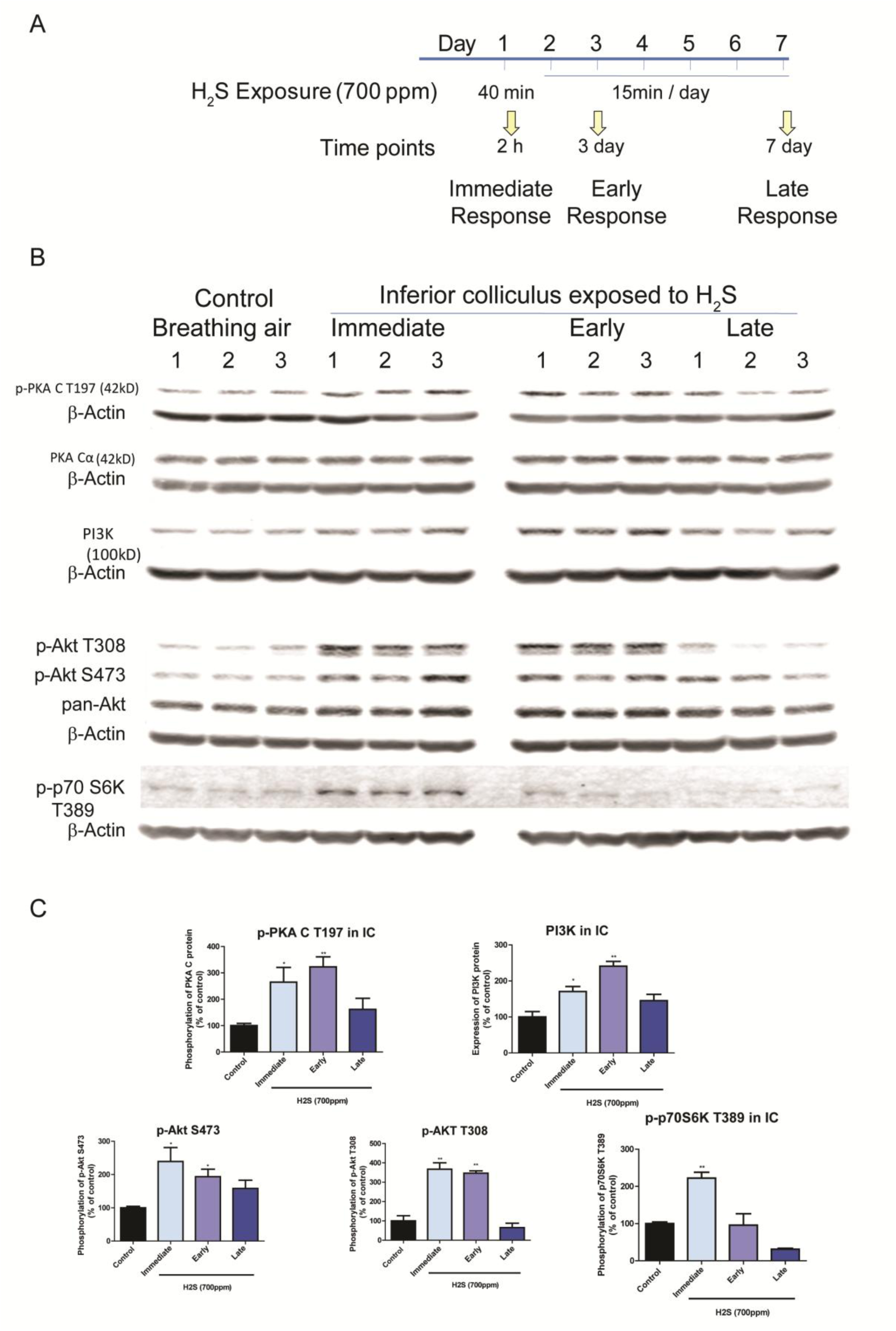
H_2_S exposure activated the PI3K/Akt signaling pathway. Activation of PI3K/Akt signaling pathways in the IC after H_2_S exposure was measured by Western blotting analyses. A, Phosphorylation of AKT at both S474 and T308 was increased at the immediate and early response time points following H_2_S exposure compared to the breathing air control group. Expression of PI3K was also upregulated at the immediate and early response times following H_2_S exposure compared to the breathing air control group. Phosphorylation of PKA C at T197 was also increased at the immediate and early response times points H_2_S exposure. Phosphorylation of p-70S6k at T389 was increased at the immediate response to H_2_S exposure. B, band intensities of Western blotting in A were measured and visualized in graph view. Significant differences were determined by t-test with comparison to the breathing air control group and were indicated with *(p<0.05) or **(p<0.01).

### 3.6 H_2_S exposure induced activation of the MAPK signaling pathway

Mitogen-activated kinases (MAPKs) transduce and amplify extracellular stimuli into a wide variety of cellular responses (Kamiyama et al. 2015). MAPKs signaling pathway also interacts with the PI3K/Akt signaling pathway (Lee et al. 2006; Pappalardo et al. 2016). Activation of MAPKs signaling pathway during acute H_2_S exposure was evaluated in the IC. Phosphorylation of stress-activated protein kinase (SAPK) was increased more than three-fold during the immediate and early phases of acute H_2_S exposure (Fig 5). Phosphorylation of p38 MAPK at Thr 180 / Tyr 182 was increased more than four-fold (Fig 5). However, ERK1/2 expressions were not altered during H_2_S exposure (data not shown). Nonetheless, the expression of the activating transcription factor 2 (ATF2), one of the downstream substrates of p38 MAPK and SAPK, was increased during the early and late phases of H_2_S exposure (Fig 5). Expression of cAMP responsive binding protein 1 (CREB-1), another downstream substrate of SAPK signaling pathway, was also increased after H_2_S exposure (Fig 5). Expression of Fos and JunB were shown to be increased at the transcription level (Fig 3).

**Figure 5.**
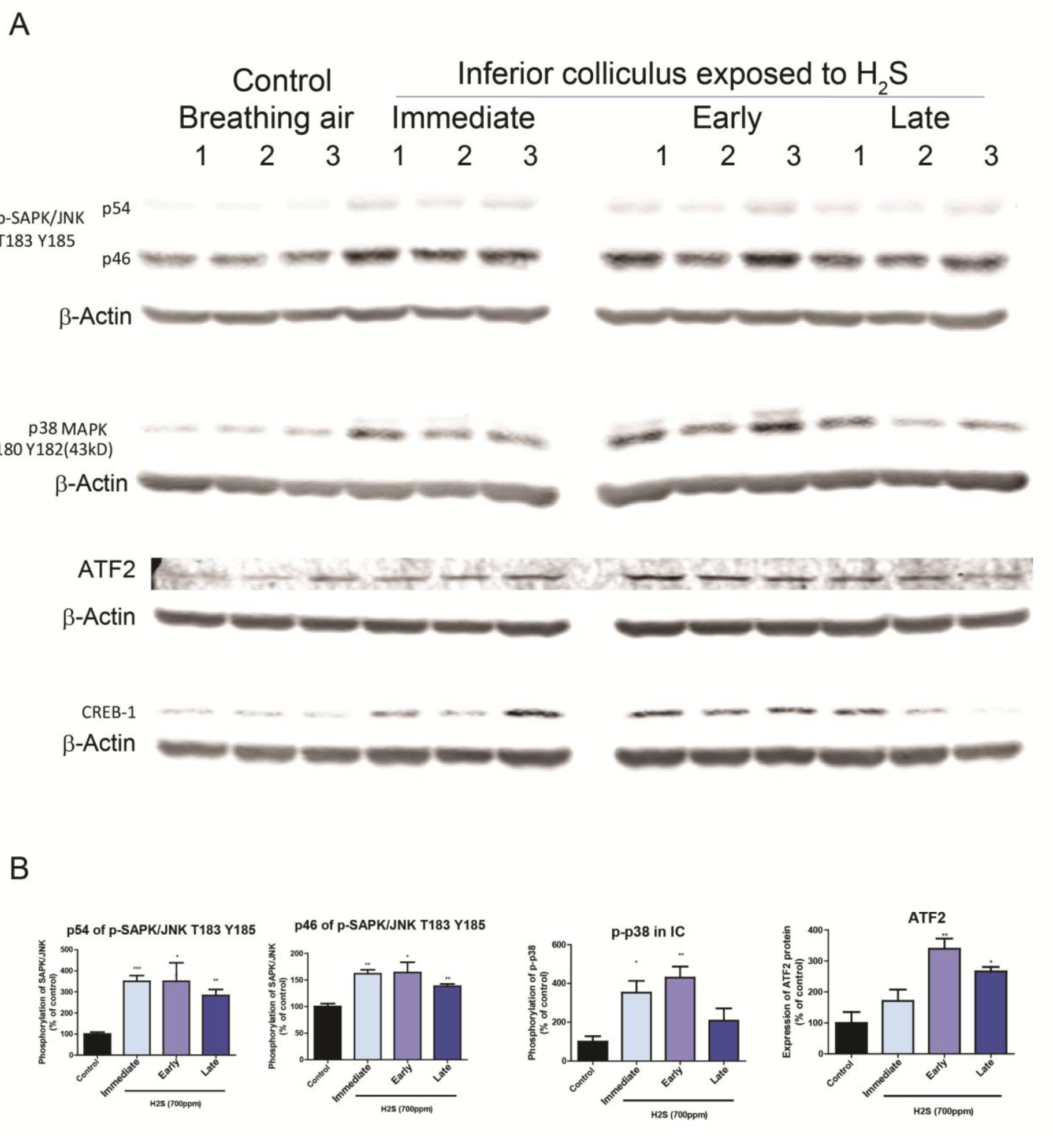
H_2_S exposure activated the MAPK signaling pathway in the IC. Activation of the MAPK signaling pathways in the IC after H_2_S exposure was measured with Western blotting analyses. A, Phosphorylation of SAPK/JNK at T183/Y185 was increased after H_2_S exposure compared to the breathing air control group. Phosphorylation of p38 was increased at the immediate and early response times following acute H_2_S exposure. Expression of ATF2 was upregulated at early and late response to H_2_S exposure compared to the breathing air control group. B, band intensities of Western blotting in A were measured and visualized in graph view. Significant differences were determined by t-test with comparison to breathing air control group and were indicated with *(p<0.05) or **(p<0.01).

### 3.7 MRI revealed injury of the IC and thalamus following acute H_2_S exposure

Previously, we reported that H_2_S exposure induced histological lesions in the IC and TH (Anantharam et al. 2017). To evaluate brain lesions following acute H_2_S exposure *in vivo*, the entire brains of live mice were scanned by coronal MRI. MRI analyses revealed hyperintense lesions in the IC and thalamus (TH) of H_2_S-exposed mice compared to breathing air control group, suggesting severe edema in these regions (Fig. 6 B). Results of lesion size are summarized in Fig. 6 C. No lesions were found in the IC and TH of control mice exposed to breathing air.

**Figure 6.**
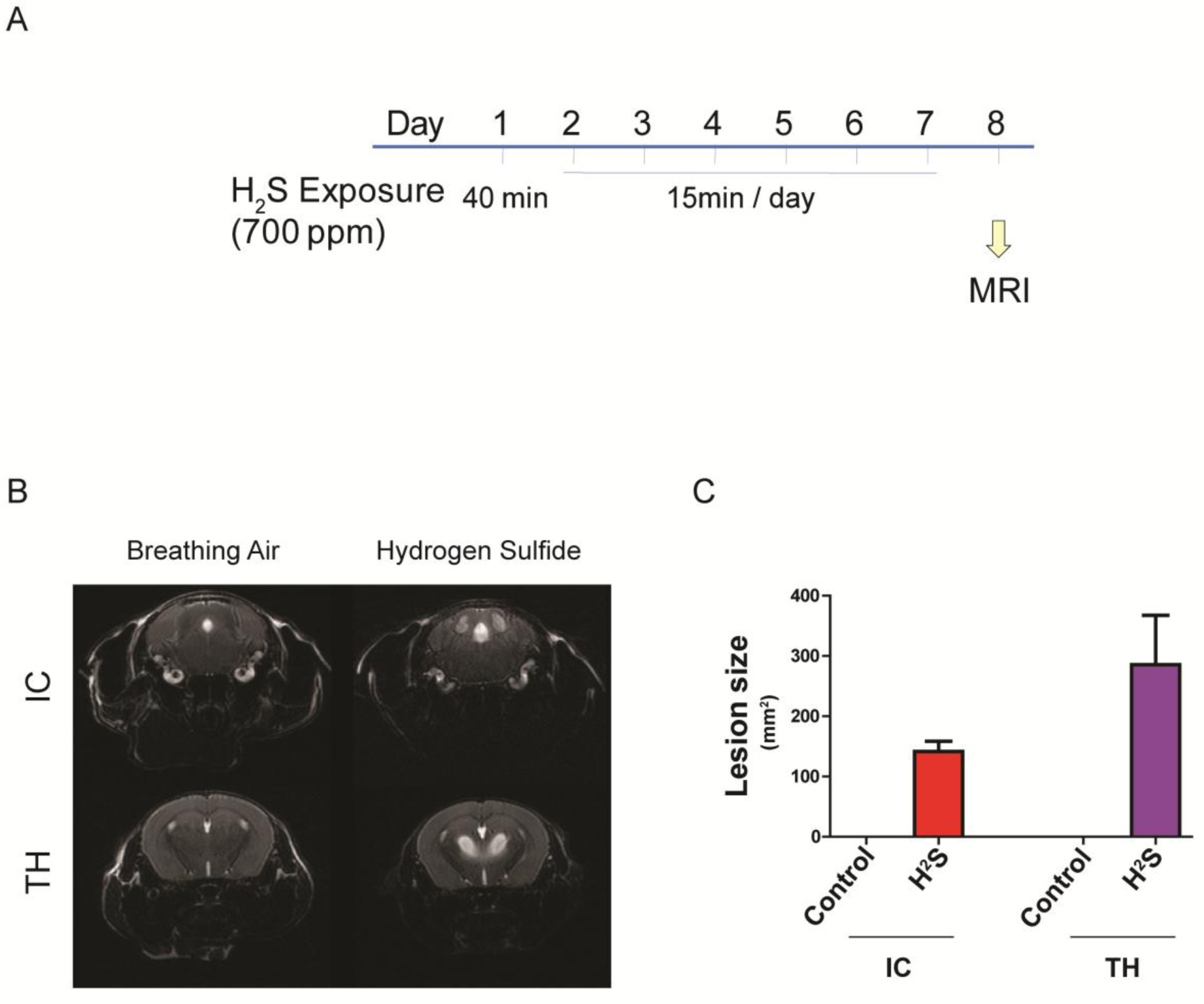
Acute exposure to H_2_S induced brain lesion in IC and TH. Mice were acutely exposed to H_2_S daily for 7 days, followed by scanning T2-weighted MRI. A scheme of the exposure paradigm is shown (A). The MRI images of the IC and TH regions are shown (B). Hydrogen sulfide exposure lesions in IC and TH were measured and visualized in graph view (C).

### 3.7 H_2_S exposure induced production of pro-inflammatory cytokines in immune cells *in vitro*

The late phase of H_2_S exposure was shown to involve many biological pathways related to inflammation. To identify the cell types involved in the inflammatory response in acute H_2_S-induced neurotoxicity, astrocyte and microglial cells were tested *in vitro*.

MMC and U373 human astrocyte cells were exposed to 1mM NaHS, a chemical H_2_S donor, for 6h. IL-1β was upregulated in both microglial and astrocyte cells, while IL-18 was upregulated in microglial cells (Fig 7).

**Figure 7.**
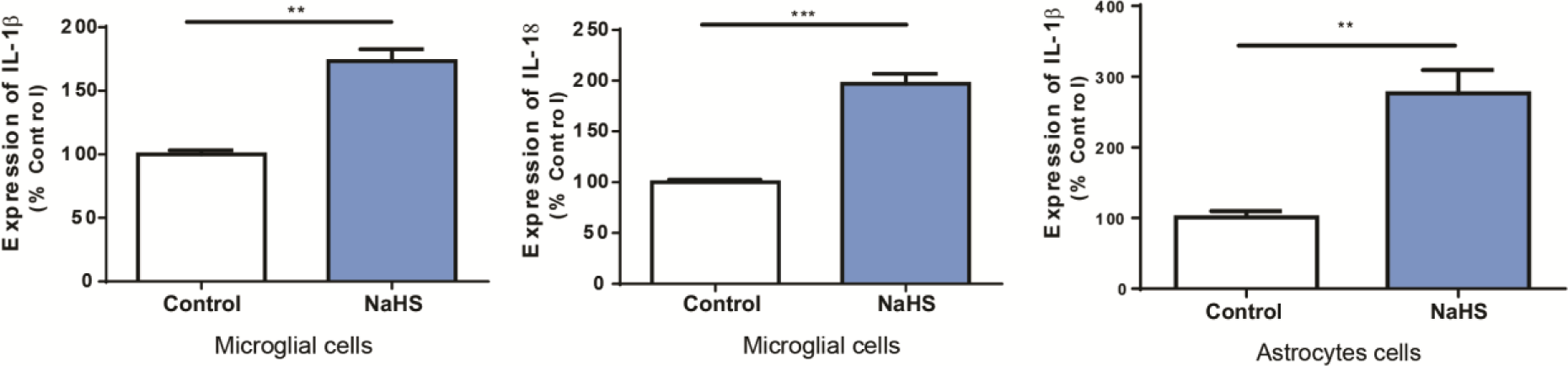
Microglia and astrocytes release cytokines following H_2_S exposure. Mouse microglial cells and human astrocyte cells were exposed to 1mM of sodium hydrosulfide (NaHS) for 6 h. Expression of IL-1β and IL-18 was examined using quantitative RT-PCR. Gene expression was compared to control group. Gapdh was used as internal reference gene. Significant differences were determined by t-test with comparison to breathing air control group and were indicated with **(p<0.01) or ***(p<0.001).

## 4. Discussion

Acute exposure to high concentration of H_2_S induces detrimental neurotoxic effects including seizures, knockdown, respiratory depression, and sudden death in animals. The exact mechanisms of H_2_S-induced neurotoxicity have not been elucidated yet, which has delayed the development of appropriate antidotes/therapies for treatment of H_2_S-induced neurotoxicity.

In this study we used the inhalational animal model of acute H_2_S toxicity developed in our lab (Anantharam et al. 2017) to assess transcriptomic analysis in the IC to identify key molecules playing important roles in the pathogenesis of H_2_S-induced neurotoxicity. A single and repeated acute exposure paradigm was used in this study, which emulates reported accidents with multiple exposure to high doses of H_2_S (Ahlborg 1951), although a single exposure is a more common accidental scenario. The exposure paradigm in this study also allows comparative effects of repeated acute exposures to a single exposure. Thus, this experimental design allowed exploration of acute H_2_S-toxicity outcomes from immediate response (same day following single acute exposure) to late response to H_2_S exposure (following repeated short acute exposures). Transcriptomic profiling analyses showed that H_2_S exposure significantly induced DEG profiling involving multiple dysregulated biological pathways, even following a single acute exposure.

One phenotype of H_2_S intoxication is seizure activity. Seizure activity can be induced by dysregulation of neurotransmitters in the central nervous system (CNS). Previous work from our laboratory has shown that acute H_2_S exposure increases brain dopamine and serotonin concentrations (Anantharam et al. 2017). In this study, catecholamine biosynthesis was suppressed as there was downregulation of dopamine beta-hydrolyxase (Dbh), dopa decaboxylase (Ddc), and tyrosine hydroxylase (Th) 2h following a single acute H_2_S exposure. The increased level of dopamine following acute H_2_S exposure may suppress the expression of genes of dopamine synthesis pathway. The transcriptomic profiling analysis also revealed that gamma-aminobutyric acid (GABA) signaling pathway was dysregulated at the immediate (single exposure) and early response phases 2 h after the second dose of acute H_2_S exposure. GABA transporters including solute carrier family 6 member 12 and solute carrier family 6 member 13 were upregulated at the immediate response phase (2 h after a single acute H_2_S exposure). Several GABA receptor subunits including GABA type A receptor alpha6 subunit and GABA type A receptor rho2 subunit were also upregulated in the early response phase. Previous work showed an increase of the glutamate:GABA ratio following H_2_S exposure (Anantharam et al. 2017). The decreased level of GABA as seen in previous studies may have induced an increase in the expression of the genes involved in GABA regulation to compensate for the deficit of GABA levels. Previously we have shown that suppression of seizure activity by treatment with an anti-convulsant drug, midazolam (MDZ), significantly rescued mice from death (Anantharam et al. 2018a). It is plausible that the effects of MDZ may have lessened the harmful effects of glutamate signaling cascade by augmenting GABA signaling cascade. Cysteine rich with EGF-like domains 2 (Creld2) may regulate transport of α4-β2 neuronal acetylcholine receptor. Creld2 was down-regulated, indicating suppression of acetylcholine signaling pathways during H_2_S exposure. These results indicate H_2_S exposure may mediate dysregulation of multiple neurotransmitter signaling pathways and this may play important roles in H_2_S-induced neurotoxicity in the IC. All these changes in neurotransmitter patterns likely contribute to H_2_S-induced seizures. More work remains to be done to piece together which of these changes play dominant roles in seizure induction.

Hydrogen sulfide is well known to induce vasodilation (Lavu et al. 2011). Endothelin1 (Edn1) works as an endothelium-derived vasoconstrictor. Edn1 was consistently upregulated more than two folds after H_2_S exposure. In addition, genes associated with activation of coagulation system were upregulated. Specifically, coagulation factor III (F3), plasminogen activator tissue type (Plat), plasminogen activator urokinase receptor (Plaur), and serpin family E member 1 (Serpine1) were all upregulated 2 h following a single acute H_2_S exposure, suggesting blood vessel injury and brain hemorrhage. F3, plat, and plaur were upregulated 2 h after a single acute H_2_S exposure and after two acute exposures (early response), while alpha-2-macroglobulin (A2m) and von Willebrand factor (Vwf) were upregulated from early response (2 exposures to H_2_S) to late response (4 exposures to H_2_S). These results further support our previous findings that H_2_S exposure may induce hemorrhage in the IC, suggesting a breach of the blood brain barrier may play a role in H_2_S-induced neurotoxicity (Kim et al. 2018a). Considering that the IC requires high blood supply for playing a role as a hub for hearing (Huffman and Henson 1990), this region has a dense blood capillary network. We postulate that H_2_S-induced blood vessel injury and subsequent hemorrhage in the IC may also contribute to hypoxic conditions in the IC.

We have previously shown that H_2_S exposure activated the hypoxic signaling pathway and upregulated Hif-1α (Kim et al. 2018a). The Akt signaling pathway has previously been shown to be activated under hypoxic conditions and reportedly plays an important role in survival of neuronal cells (Alvarez-Tejado et al. 2001); it also mediates the Hif-1α signaling pathway (Kilic-Eren et al. 2013). Transcriptomic analysis revealed H_2_S exposure induced activation of Akt signaling pathway at the immediate and early phases. Concomitantly, PI3K was significantly upregulated at the immediate and early response phases. In addition, PKA was also activated at the immediate and early phases following acute H_2_S exposure. p70 S6K, one of substrates of Akt signaling pathway, was also activated immediately at the immediate phase following acute H_2_S exposure. Collectively, this data indicates that H_2_S exposure induced hypoxic conditions in the IC which then activated the Akt pathway for the survival of neuronal cells in the IC.

This study also provided more evidence on the involvement of oxidative stress in the pathogenesis of acute H_2_S-exposure. Hydrogen sulfide exposure consistently induced oxidative stress response in the IC throughout the course of this study, starting from 2 h following a single acute exposure (immediate) to following 4 exposures (late response). Impairment of cytochrome c oxidase by ethanol overdose was previously reported to generate ROS (Srinivasan and Avadhani 2012). Hydrogen sulfide may generate ROS by inhibiting cytochrome c oxidase in a similar manner. In our previous studies we have demonstrated that H_2_S-inhibits cytochrome c oxidase (Anantharam et al. 2017); moreover, H_2_S exposure also led to the generation of hydroxyl radicals (Jiang et al. 2016) all of which contribute to oxidative stress. In our previous work, we have reported that H_2_S exposure activated the Nrf2 signaling pathway (Kim et al. 2018a), further supporting the hypothesis of H_2_S-induced oxidative stress as a toxic mechanism. MAF BZIP transcription factor F (Maff) is a basic leucine zipper transcription factor and forms a heterodimer with Nrf2 to induce antioxidant or electrophile response elements (AREs) (Katsuoka et al. 2005). In this study we have shown that H_2_S exposure consistently induced upregulation of Maff and Mafk following H_2_S exposure. Uncoupling protein 2 (Ucp2) is a mitochondrial transporter protein and was shown to be upregulated 70 - 100% after H_2_S exposure in the IC compared to breathing air control group. Ucp2 creates proton leakage across the inner mitochondrial membrane to uncouple oxidative phosphorylation from ATP synthesis, which plays a role in controlling ROS production (Mailloux and Harper 2011) and in calcium regulation (Motloch et al. 2016). Collectively, these data further show that H_2_S exposure induces significant oxidative stress. Thus therapies targeting inhibition of H_2_S-induced oxidative stress may be good candidates for treating H_2_S-induced neurotoxicity.

Hydrogen sulfide exposure induced dysregulation of the calcium signaling pathway in this study, indicating intracellular calcium may be dysregulated. It is reported that H_2_S exposure increased intracellular Ca^2+^ concentration via PKA and PLC/PKC pathways (Yong et al. 2010). Hydrogen sulfide exposure indeed activated PKA by phosphorylation at T197 in this study. Multiple factors such as impaired cytochrome c oxidase activity with subsequent reduced energy production, oxidative stress, and Ca^2+^ dysregulation may collectively induce endoplasmic reticulum (ER) stress, leading to unfolded protein response which we observed in this study. Nonetheless, further research is required to examine how H_2_S exposure exactly induces the unfolded protein response. However, RNA Seq analyses in this study revealed several heat shock protein family genes were downregulated after H_2_S exposure including HSPA1b, HSPA2, HSPA5, HSPA1A. HSPA5 encodes the binding immunoglobulin protein which serves an important role in ER stress (Wang et al. 2017). Hydrogen sulfide exposure also downregulated X-Box binding protein 1 (Xbp1). Xbp1 plays an important role in unfolded protein response (UPR) (Bahar et al. 2016). Interestingly, Mitogen-activated protein kinase 6 (Map3k6) was upregulated in the IC of H_2_S exposed mice more than two to three folds. Apoptosis signaling kinase 1 (Ask1) responds to ER stress and induces JNK activation (Hattori et al. 2009). Map3k6 functions to activate Ask1 by phosphorylation. Indeed, acute H_2_S exposure activated JNK and p38 signaling pathways. Several Jun proteins including Jun proto-oncogene (Jun), Jun B proto-oncogene (JunB), and jun D proto-oncogene (JunD) were also consistently upregulated after H_2_S exposure following either a single exposure (immediate), two exposures (early), and 4 exposures (late). Collectively, this data implicates the endoplasmic stress and unfolded protein response as a potential key mechanism leading to cell death following exposure to acute H_2_S poisoning. It may be worthy to explore targeting this pathway as a potential therapeutic approach for treatment of acute H_2_S-induced neurotoxicity.

The cell death signaling pathway was dysregulated 2 h following a single acute exposure to H_2_S. For example, expression of the anti-apoptotic gene, Bcl-2, was downregulated 2 h after a single H_2_S exposure. p53 signaling pathway which senses DNA damage was also activated at 2 h following a single acute H_2_S exposure. Cyclin D2 (Ccnd2), cycline-dependent kinase inhibitor 1A (Cdkn1a), growth arrest and DNA damage inducible gamma (Gadd45g), tumor protein p53 inducible nuclear protein 1 (Trp53inp1), and Fas were also upregulated 2 h following a single acute exposure to H_2_S (immediate response), indicating early activation of the p53 signaling pathway. Fat atypical cadherin 2 (Fat2) is a tumor suppressor gene which plays a role in controlling cell proliferation (Katoh 2012). Expression of Fat2 was consistently upregulated following H_2_S exposure indicating that cell death signaling was activated. Cd93 is a cell-surface glycoprotein and was shown to be involved in intercellular adhesion and in the clearance of apoptotic cells. Consistent upregulation of Cd93 following H_2_S exposure may indicate that H_2_S exposure induced cell death during H_2_S exposure. However, NFkB inhibitor, alpha (Nfkbia), was consistently upregulated, while Tumor Necrosis Factor (Ligand) Superfamily, Member 10 (Tnfsf10) was consistently downregulated. Collectively, several cell survival and cell death signaling pathways were involved in H_2_S exposure response. The meaning of all these changes is not clear from this one study. Clearly, a better understanding of cell survival and death signaling pathways in detail requires further studies.

To gain insight of brain pathology in live H_2_S exposed mice, non-invasive MRI scanning imaging system was used. T2 weighted imaging showed edema in the IC and TH of mice indicating significant neurodegeneration in these two brain regions on day 8 following H_2_S exposure. These MRI results are consistent with our previous histology and immunohistochemistry results (Anantharam et al. 2017; Kim et al. 2018a) which showed that H_2_S exposure induced severe loss of neurons and activation of neuroinflammation starting on day 3. Both the IC and TH have been linked to seizures. The origin of H_2_S-induced seizure is yet to be determined. The IC is a part of the brain stem which serves as a major auditory center in the brain. Interestingly, the IC has been linked to audiogenic seizure (Garcia-Cairasco et al. 1996). GABA plays an important inhibitory role whereas glutamate-mediated stimulation is excitatory to neurons in the IC (Faingold 2002). The TH has also been linked to different forms of seizures including absence seizure, temporal lobe seizures and generalized seizures (Carney and Jackson 2014). We previously showed that pre-treatment of mice with MDZ, an anticonvulsant agent, protected mice from H_2_S-induced seizures and death (Anantharam et al. 2018b). Midazolam pretreatment also significantly reduced lesion severity in these two regions. This data suggests that seizures play important roles in H_2_S-induced neurotoxicity, cell death, and evolution of neurological sequalae. In other words, seizures may be linked to the development of lesions both in the IC and the TH since pretreatment with midazolam significantly reduces lesion severity in both regions.

The analysis for potential upstream regulator(s) after H_2_S exposure revealed an inflammatory response (e.g. TNFα, IL-1β, and IL-6) in addition to transcription factors for cell proliferation and death, such as p53, in all phases of H_2_S-induced neurotoxicity examined in this study. These data indicate that the H_2_S exposure immediately induces a pro-inflammatory response as IL-1β was upregulated immediately following acute H_2_S exposure. IL-1β, IL-18, IL-6, Ccl2, and TNF-α were all upregulated in IC during the early and late phases after H_2_S exposure. *In vitro* cell culture studies further showed that mouse microglial cells produce IL-1β and IL-18, while human astrocytes cells produce IL-18 at 6h in response to H_2_S exposure. Previously we have shown that glial fibrillary acidic protein (GFAP) was upregulated by acute H_2_S exposure (Anantharam et al. 2017). Collectively, these data show that microglia and astrocytes immediately respond to H_2_S exposure and elicit initial neuro-inflammatory reactions. In turn, these reactions unleash powerful neuroinflammation with production of Ccl2, TNF-α, and IL-6 leading to neuronal loss and neurodegeneration in the IC (Anantharam et al. 2017). The implication of these results is that drugs which suppress neuro-inflammation may be potential candidates for counteracting H_2_S-induced neuroinflammation.

## 5. Conclusions

In the past, the mechanisms of H_2_S-induced neurotoxicity and neurodegeneration have remained poorly understood. In this study, we performed transcriptomic analysis of the IC in mice following acute H_2_S exposure. Results revealed important key molecules and molecular pathways including unfolded protein response, neurotransmitters, calcium dysregulation, oxidative stress, survival/death, hypoxia, and inflammatory response in the IC following acute H_2_S exposure. Potential upstream regulators in IC were found to be inflammatory initiators such as TNF, IL-1β and IL-18. These results are consistent with some of the previous findings which we have published. This study has also proved that *in vivo* MRI analysis is effective in imaging the lesions in IC and TH following H_2_S exposure. *In vitro* studies showed that microglia and astrocytes released IL-1β and IL-18 which we hypothesized to be key molecules in orchestrating multiple biological pathways. The overall hypothetical schemes of key events are presented in Figs 8 and 9. Overall, this novel study has shown that multiple complex biological pathways are involved in the pathogenesis of H_2_S-induced neurotoxicity. The common simplistic dogma that H_2_S-induced neurotoxicity is caused by inhibition of cytochrome c oxidase leading to energy deficit and cell death should be critically questioned and explored while also examining other mechanistic pathways. More research is warranted to further define the exact molecular mechanisms involved in H_2_S-induced neurotoxicity. Collectively, this study has identified important key elements, biological pathways, and potential signal cascade initiators involved in H_2_S-induced neurotoxicity and the evolution of neurological sequelae. More in-depth research is needed to further define the pathways leading to neuronal cell death and more definitive therapeutic targets.

**Figure 8.**
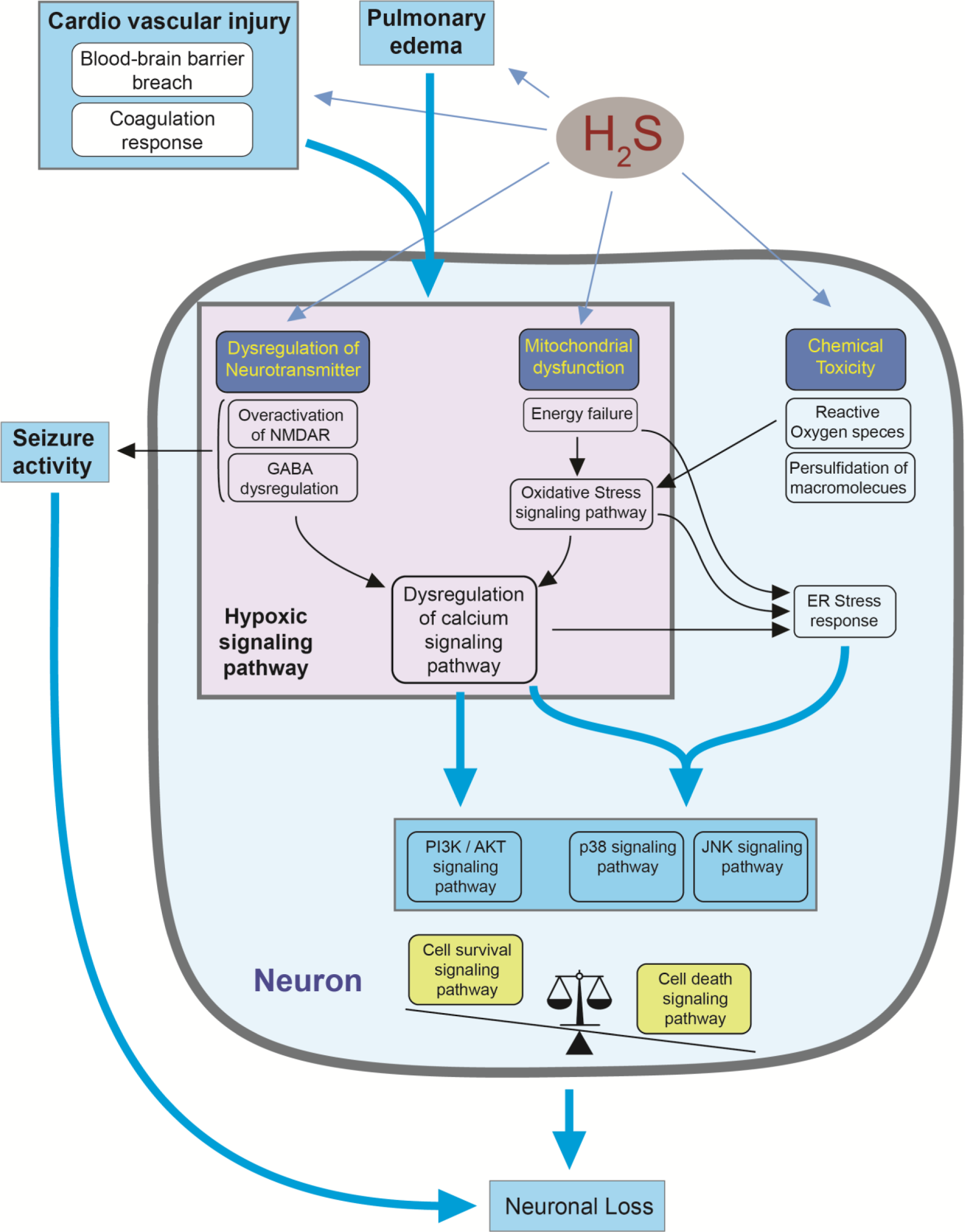
A hypothetical diagram summarizing the pathways leading to H_2_S-induced neuronal cell death. Hydrogen sulfide exposure induces dysregulation of multiple biological pathways leading to neuronal loss in the IC. Pulmonary edema and cardiovascular injury may reduce supply of oxygen, which may play a role in inducing the hypoxic signaling pathway. In addition, dysregulation of neurotransmitters such as NMDA and GABA and mitochondrial dysfunction may induce hypoxic signaling pathway, which may lead to calcium dysregulation and cell survival and death signaling pathways. Mitochondrial dysfunction and dysregulated calcium signaling may induce ER stress response and cell death signaling pathway. When cell death signaling pathway overcomes survival signaling pathway, neuronal loss may occur. Seizure activity derived from dysregulation of neurotransmitter system may further affect on neuronal loss.

**Figure 9.**
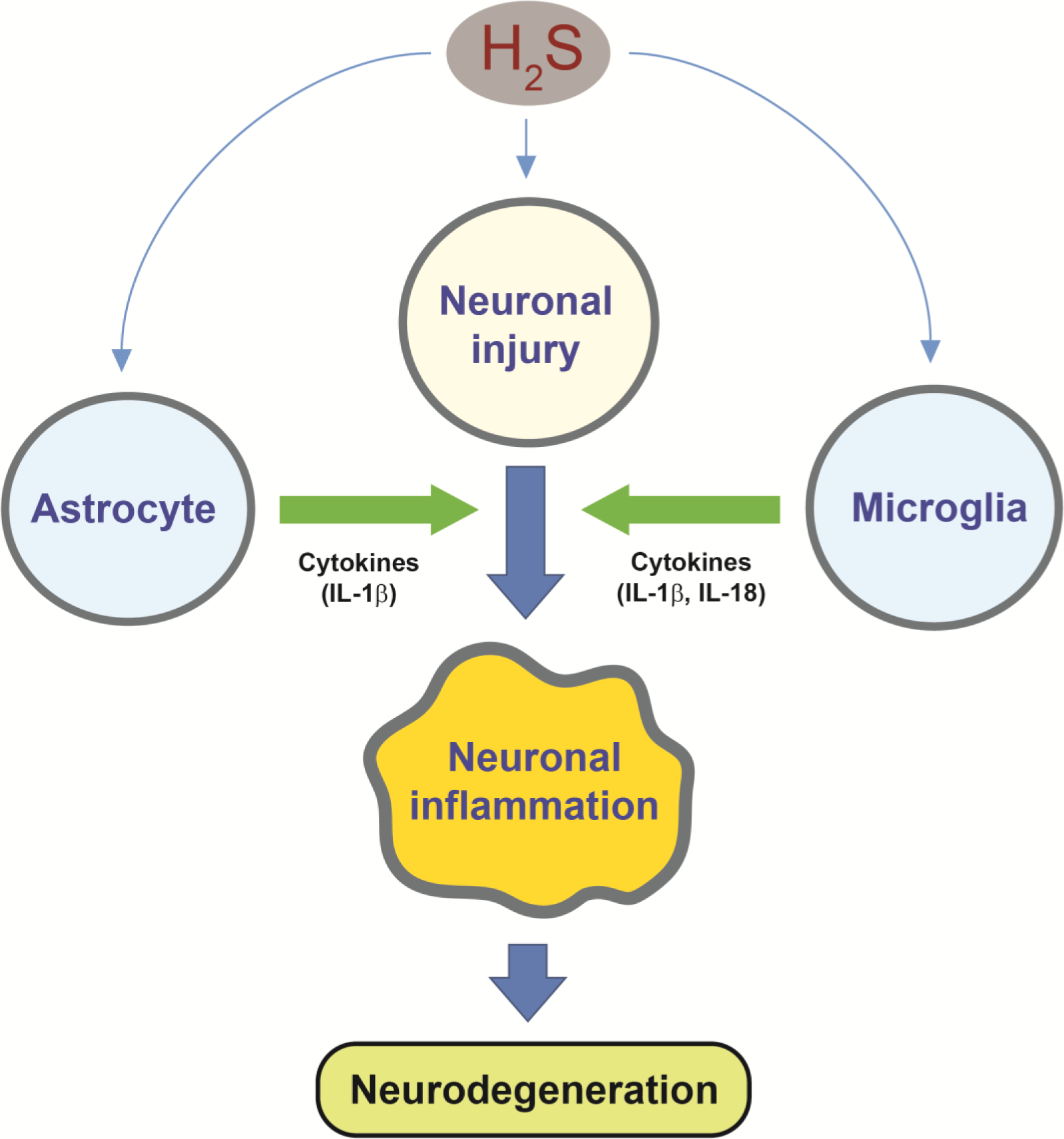
H_2_S exposure activates astrocyte and microglia in addition to neuronal injury. Pro-inflammatory cytokines such as IL-1β and IL-18 from astrocytes and microglia may accelerate neuronal inflammation leading to massive neurodegeneration in IC.

## Acknowledgment

The authors do not have any conflict of interest.

This work was supported by Iowa State University College of Veterinary Medicine Seed grant and Iowa State University Tuskegee Collaboration Seed grant. None of the funder played a role in any of the studies presented in this manuscript.

